# The Insulin receptor regulates the persistence of mechanical nociceptive sensitization in flies and mice

**DOI:** 10.1101/2022.04.02.486841

**Authors:** Yan Wang, Roger Lopez-Bellido, Xiaojiao Huo, Annemieke Kavelaars, Michael J. Galko

## Abstract

Early phase diabetes is often accompanied by pain sensitization. In the fruit fly *Drosophila*, the insulin receptor (InR) regulates the persistence of injury-induced thermal nociceptive sensitization. Whether *Drosophila* InR also regulates the persistence of mechanical nociceptive sensitization remains unclear. Mice with a sensory neuron deletion of the gene encoding the Insulin receptor (*Insr*) show normal nociceptive baselines, however, it is not known whether deletion of *Insr* in nociceptive sensory neurons leads to persistent nociceptive hypersensitivity in an inflammatory pain paradigm. In this study, we used fly and mouse nociceptive sensitization models to address these questions. In flies, *InR* mutants and larvae with sensory neuron-specific expression of RNAi transgenes targeting *InR* exhibited persistent mechanical hypersensitivity, as previously observed for the thermal sensory modality. Mice with a specific deletion of the *Insr* gene in NaV1.8+ nociceptive sensory neurons showed normal nociceptive thermal and mechanical baselines similar to controls. In an inflammatory paradigm, however, these mutant mice showed persistent mechanical (but not thermal) hypersensitivity, particularly in female mice. Mice with the NaV1.8+ sensory neuron specific deletion of *Insr* did not show metabolic abnormalities that would be typical of a systemic defect in insulin signaling. Our results show that some aspects of the regulation of nociceptive hypersensitivity by the Insulin receptor are shared between flies and mice and that this regulation is likely independent of metabolic effects.

## Introduction

The fruit fly (*Drosophila melanogaster*) has emerged over the last decade as a powerful system to genetically dissect nociception and nociceptive sensitization (Im and Galko 2012; Milinkeviciute et al. 2012). Early work established that *Drosophila* avoid noxious stimuli through conserved transient receptor potential (TRP) channels (Hwang et al. 2012; Kang et al. 2010; Tracey et al. 2003). These avoidance responses, such as the aversive rolling of *Drosophila* larvae, are enabled by multidendritic peripheral sensory neurons (Gao et al. 1999; Hwang et al. 2007) whose elaborate dendritic arbors tile over the barrier epidermis. These neurons are the functional counterparts of unmyelinated nociceptors in vertebrates. Nociceptive responses in *Drosophila* include detection of noxious heat (Babcock et al. 2009; Tracey et al. 2003), cold (Turner et al. 2016), mechanical (Kim et al. 2012; Lopez-Bellido et al. 2019b; Mauthner et al. 2014), and chemical (Lopez-Bellido et al. 2019a) stimuli. There is additional complexity to the response, however. As in vertebrates (Gold and Gebhart 2010), tissue injury is capable of causing transient hypersensitivity to both noxious (hyperalgesia) and non-noxious (allodynia) stimuli. Some of the molecular pathways that mediate these responses such as Tumor Necrosis Factor (TNF) signaling (Babcock et al. 2009; Cunha et al. 1992; Cunha et al. 2005), Substance P/Tachykinin (Im et al. 2015; Laird et al. 2001; Weng et al. 2001), and Hedgehog signaling (Babcock et al. 2011; Liu et al. 2018) are conserved. Finally, flies possess second-order and higher order central interneurons (Hu et al. 2017; Ohyama et al. 2015; Yoshino et al. 2017) that comprise a nociceptive circuit and confer the potential for nociceptive responses to be modulated by competing sensory inputs.

Flies have also proven to be a useful model of insulin-like signaling (ILS). *Drosophila* possess a clear insulin receptor ortholog, InR (Fernandez et al. 1995), insulin-like peptides (Rulifson et al. 2002), and conserved downstream pathway components (Clancy et al. 2001; Goberdhan et al. 1999). Disruption of ILS through mutation or *in vivo* RNAi-mediated knockdown leads to a wide variety of phenotypes including increased longevity (Tatar et al. 2001), decreased cell size (Böhni et al. 1999; Rulifson et al. 2002), and metabolic effects that mirror aspects of diabetes in vertebrates (Musselman et al. 2011). A prior study from our lab indicated that *Drosophila* InR is required specifically in nociceptive sensory neurons for the resolution of injury-induced thermal nociceptive sensitization (Im et al. 2018). In this same study (Im et al. 2018), persistent injury-induced nociceptive hypersensitivity was also observed in *Drosophila* larvae mimicking type I diabetes (Rulifson et al. 2002) or type II diabetes (reared on a high sugar diet) (Musselman et al. 2011), suggesting a potential connection between sensory neuron loss of ILS and hypersensitivity. Here, we test whether fly InR is similarly required for the persistence of mechanical nociceptive hypersensitivity.

In mice, homozygous *Insr* mutants are lethal (Accili et al. 1996). Tissue specific deletion of *Insr* combining a floxed allele (Bruning et al. 1998) with a Nestin-Cre driver that expresses in neuronal and glial precursors led to no obvious sensory phenotypes (Bruning et al. 2000). As with most tissues, *Insr* is reported to be expressed in sensory neurons (Sugimoto et al. 2002; Sugimoto et al. 2000). A more specific deletion of *Insr* in sensory neurons using *Advillin-Cre* (Zurborg et al. 2011), revealed no obvious alterations to baseline nociceptive sensitivity in the absence of injury (Grote et al. 2018). In this latter study, systemic impacts on sugar metabolism (elevated serum insulin and glucose intolerance) were observed (Grote et al. 2018). Here, motivated in part by our prior studies in *Drosophila*, we tested whether deletion of *Insr* in NaV1.8-positive nociceptive sensory neurons (Agarwal et al. 2004), regulates the onset or duration of thermal or mechanical hypersensitivity or impacts systemic measures of metabolism.

## Materials and Methods

### Fly Genetics

*Drosophila* were reared on standard cornmeal medium under a 12 h light-dark cycle. All crosses were cultured at 25 °C except for the *InR* transheterozygotic combination (*InR^e19/93Dj4^*) (Tatar et al. 2001), which was reared at 18 °C until the third larval instar and then moved to 25 °C for experiments. *w^1118^* and/or *Gal4^109(2)80^/+* (referred to as *md-Gal4*) crossed to *w^1118^* served as control strains. *InR* mutant alleles used were a transheterozygous combination of *InR^e19^* and *InR^93Dj4^*. Both are loss of function measured by a decrease of kinase activity (Chen et al. 1996; Tatar et al. 2001). Tissue-specific expression of *UAS* transgenes was controlled by *md-Gal4* (Gao et al. 1999) which expresses in all four classes of multidendritic (md) sensory neurons. The *UAS-RNAi* lines (Ni et al. 2011) used was *InR^JF01482^*.

### Mouse Genetics

All experimental protocols were approved by the Institutional Animal Care and Use Committee (IACUC) at The University of Texas MD Anderson Cancer Center and conformed to the U.S. National Institutes of Health guidelines on the ethical care and use of animals. Mice were housed 1–5 per cage and maintained on a 12-h light/dark schedule in a temperature-controlled environment with free access to food and RO water.

*Scn10a^tm2(cre)Jwo^* (*NaV1.8^Cre/+^*) mice have been previously characterized, demonstrating sensory neuron specific Cre recombinase activity (Agarwal et al. 2004). *Insr^tm1Khn^ Insr^lox/lox^*) mice with *loxP* sites flanking exon 4 of the insulin receptor gene (Bruning et al. 1998) were purchased from the Jackson Laboratory (JAX stock #006955, Bar Harbor, ME). Cre-mediated deletion of exon 4 would create a nonfunctional 308 amino acid truncated protein. *NaV1.8^Cre/+^* mice were bred to *Insr^lox/lox^* to generate heterozygous *NaV1.8^Cre/+^, Insr^lox/+^* mice. *NaV1.8^Cre/+^, Insr^lox/+^* were bred to *Insr^lox/lox^* to produce mice with an *NaV1.8^Cre/+^, Insr^lox/lox^* genotype and littermate controls. These mice should have a specific deletion of Insr exon 4 within NaV1.8-expressing nociceptive sensory neurons. Mice were genotyped from DNA harvested from ear punches by Transnetyx (Cordova, TN).

### *Drosophila* mechanical nociceptive sensitization assay

To create a UV-induced tissue injury (Babcock et al. 2009), mid L3 larvae anesthetized with anhydrous ethyl ether (Thermo Fisher Scientific, Waltham, MA) were mounted on microscope slides so that the dorsal side was exposed to UV or mock-treated with 18-20 mJ/cm^2^ (measured with a UV spectrophotometer [Accumax XS-254; Spectronics Corporation, Melville, NY]) in a Spectrolinker XL-1000 ultraviolet crosslinker (Spectronics Corporation, Melville, NY). The mechanical nociception assay for *Drosophila* larvae was performed as described in (Lopez-Bellido et al. 2019b). Briefly, custom made mechanical probes exerting mechanical pressures of 200 kPa (to measure mechanical allodynia) or 462 kPa (to measure. mechanical hyperalgesia) were applied onto the posterior dorsal side of the larva at approximately abdominal segment A8 until the probe bends and exerts a constant pressure. The percentage of larvae that exhibited a nocifensive response– defined as a complete roll of 360° along the long axis of its body (within 2 s of probe bending) was recorded.

### Mouse nociception behavioral assays

#### Von Frey Test

Mice of 10-18 weeks old were acclimated to the mesh grid in individual plastic animal enclosures of 10 x 10 x 13 cm (IITC Life Sciences, Woodland Hills, CA). Assay performer was blind to the genotype of mice. The up-down method (Dixon 1980) was used to test mechanical sensitivity of the hindpaws. A set of von Frey monofilaments (Exacta Touch Test Sensory Evaluator, North Coast Medical, Gilroy, CA) that exert forces of 0.008, 0.02, 0.04, 0.07, 0.16, 0.4, 0.6, 1, 1.4, and 2 g were applied to the plantar surface of the hind paw of each mouse, starting with the 0.4 g filament (Dixon 1980). The duration of each stimulus was approximately 1 s and the inter-stimulus interval was approximately 30–60 s. The 50% withdrawal threshold for each hind paw was then calculated (Chaplan et al. 1994) with a customized Python script. The 50% paw withdrawal threshold for each mouse was averaged from both hind paws. Baseline paw withdrawal threshold was averaged from measurement on three different days.

#### Hargreaves Test

Mice of 10-13 weeks old were acclimated to the glass surface in individual plastic animal enclosures (IITC Life Sciences, Woodland Hills, CA) as per an established protocol (Hargreaves et al. 1988) with modifications (Cheah et al. 2017). Assay performer was blind to the genotype of mice. The glass surface was maintained at 30°C. A focused radiant heat light beam of 30% intensity was chosen to elicit an average baseline withdrawal latency (sudden paw lifting from the glass) of ~ 7 s and was aimed at the plantar surface of the hind paw and the paw withdrawal latency was recorded, repeating the procedure 5 times. The inter-test interval was at least 3 minutes. The average of three measurements after removing the minimal and maximal values (Cheah et al. 2017) was calculated for each hind paw with a customized Python script. The average paw withdrawal latency for each mouse was calculated from both hind paws. Baseline paw withdrawal latency was averaged from measurement on three different days prior to sensitization (below).

#### Inflammatory sensitization

Local inflammation in the paw was induced by intraplantar administration of Complete Freund’s Adjuvant (CFA, 5 μL/hind paw; Sigma-Aldrich, St. Louis, MO). The average paw withdrawal latency for mechanical and thermal nociception (see assays above) was measured on days 1, 3, 7, 10, 14 or 15, 21 after CFA administration.

### Mouse metabolism assays

Metabolic parameters for both control mice and and mice with sensory-specific deletion of *Insr* were monitored at 6, 9, 13, 17, and 21 weeks of age. Mice weights were measured at 4, 8, 12, 16, and 20 weeks of age.

#### Glucose and Hemoglobin A1C

Blood glucose levels were determined using an Ascensia Contour Next Ez Blood Glucose Monitoring System (Parsippany, NJ). Mice were fasted 3 hours prior to glucose measurement and blood was collected via tail snip at 6, 9, 13, 17, and 21 weeks of age. In addition, long term glucose levels were assessed by determining hemoglobin A1C levels at 21 weeks of age using A1CNow+ Professional Multi-Test HbA1c System (Pts Diagnostics, Indianapolis, IN).

#### Intraperitoneal Glucose Tolerance Test (IPGTT)

At 22 weeks of age, glucose tolerance was analyzed with an intraperitoneal glucose tolerance test (IPGTT) for both *Insr^lox/lox^* controls and *NaV1.8^Cre/+^*; *Insr^lox/lox^* experimental mice. After a 5-hour fast, mice were administered 20% dextrose in saline at 10 ml/kg body weight via intraperitoneal injection. Blood glucose measurements were taken immediately prior to glucose administration and at 15, 30, 60 and 120 minutes thereafter.

### *Insr* deletion and expression analysis

#### PCR

Genomic DNA was isolated from the dorsal root ganglion (DRG) and hindpaw skin of controls and *NaV1.8^Cre/+^*; *Insr^lox/lox^* mice to assess *Cre/loxP* recombination and deletion of *Insr* exon 4. The *NaV1.8-Cre* allele was detected using primers 5’-TGTAGATGGACTGCAGAGGATGGA-3’ and 5’-AAATGTTGCTGGATATTTTTACTGCC-3’ as described in (Gautron et al. 2011). The *Insr^lox/lox^* allele was detected using primers 5’-GATGTGCACCCCATGTCTG-3’ and 5’-CTGAATAGCTGAGACCACAG-3’ as described in (Bruning et al. 1998). PCR conditions were 94°C for 4 min followed by 30 cycles of 94°C for 40 secs, 60°C for 30 secs, 72°C for 25 secs and a final extension at 72°C for 8 min. Cre/lox recombination and deletion of the exon 4 of the *Insr* gene was detected using primers 5’-ATGGTGGCATGCACTTATGA-3’ and 5’-TGCCTCAGCCTCCTGAATAG-3’. PCR conditions were 94°C for 4 min followed by 35 cycles of 94°C for 40 secs, 54°C for 30 secs, 72°C for 25 secs and a final extension at 72°C for 8 min.

#### Immunofluorescence, microscopy and quantification of IRα

DRGs were dissected and fixed in 4% paraformaldehyde over night at 4°C followed by cryoprotection in 30% sucrose over night at 4°C. Fixed samples were embedded in O.C.T (Thermo Fisher Scientific, Waltham, MA). for cryosectioning at 10 μm with a Cryostar NX70 cryostat (Thermo Scientific, Waltham, MA). Collected sections were mounted on glass slides, blocked in blocking buffer (5% goat serum, 0.3% Triton X-100 in PBS) (all materials here from Thermo Fisher Scientific, Waltham, MA except goat serum-Sigma-Aldrich, St. Louis, MO) for 1 hr, and stained with primary antibodies overnight at 4 °C. The primary antibodies were guinea pig anti-NaV1.8 (1:200 for Fig. 2B; 1:100 for Fig. 2C; AGP-029, Alomone Labs, Jerusalem, Israel), chicken-anti-GFAP (1:100, PA1-10004, Thermo Fisher Scientific, Waltham, MA), and rabbit anti-IRa (1:50 for Fig. 2B; 1:25 for Fig. 2C; ab1500 Abcam, Cambridge, MA). Dilutions were in in antibody dilution buffer (1% BSA, 0.3%Triton X-100 in PBS-all materials here from Thermo Fisher Scientific, Waltham, MA). After washing with wash buffer (0.05% Tween-20 in PBS), the samples were incubated in secondary antibodies for 1 hr at room temperature. Secondary antibodies were 1:200 goat anti-guinea pig Alexa Fluro 488 (1:200 for Fig. 2B; 106-545-003, Jackson ImmunoResearch, West Grove, PA), goat anti-guinea pig Cy3 (1:25 for Fig. 2C; 106-165-003, Jackson ImmunoResearch, West Grove, PA), goat anti-chicken Alexa Fluor 488 (1:100; A11039, Thermo Fisher Scientific, Waltham, MA), and goat anti-rabbit Alexa Fluor 647 (1:200 for Fig. 2B and 1:100 for Fig. 2C; ab150083, Abcam, Cambridge, MA). After washing, the sections were coverslipped with Vectashield (H-1000, Vector Laboratories, Burlingame, CA) for microscopy. Images were captured with an Olympus FV 1000 laser scanning confocal microscope and a PlanApo N (60x/1.42 Oil) objective using FLUOVIEW version FV10-ASW 3.1 software and processed with ImageJ. The images used for quantification of the intensity of IRα were captured with a Leica MZ16 FA fluorescent stereomicroscope equipped with a PLAN APO 1.6x stereo-objective and a HAMAMATSU ORCA-ER C-4742-80-12AG digital camera using Leica LAS X. The mean intensity of IRα in NaV1.8+ and GFAP+ DRG cells was measured using Image J.

### Statistical analyses

Mouse behavioral data: All data is reported as the mean ± standard error of the mean. To compare two groups without time factor, two-tailed unpaired Student’s t test was used. To compare two groups with time factor, two-way repeated measures ANOVA followed by Sidak’s multiple comparison test was used. To compare more than two groups with time factor, two-way repeated measures ANOVA followed by Tukey’s multiple comparison test was used. Male and female mice were initially analyzed separately and the data was grouped if no sex differences were apparent. Graphpad Prism software v8 (GraphPad Software, La Jolla, CA, USA) was used for all statistical analyses (including for the fly data, as described below).

Fly behavioral data: All the data were tested for a normal distribution using Kolmogorov–Smirnov (KS) or Shapiro–Wilk normality test. To compare two groups, two-tailed unpaired Student’s t test was used. To compare more than two groups, a one-way ANOVA (followed by Tukey post hoc test) was used.

For all behavioral data: Asterisks in the graphs indicate the significance of p-values comparing indicated groups: *p < 0.05, **p < 0.01, ***p < 0.001.

## Results

### *Drosophila InR* regulates the persistence of mechanical nociceptive sensitization

We first set to test whether the persistent thermal nociceptive sensitization observed in flies with *InR* mutation and with sensory neuron-specific expression of *UAS-InR^RNAi^* transgenes (Im et al. 2018) extends to mechanical nociception. To do this, we used a newly refined assay for mechanical nociception (Lopez-Bellido and Galko 2020; Lopez-Bellido et al. 2019b) (see schematic, Figure 1A). In the absence of injury, control larvae showed a normal dose-response curve to sub-noxious (≤ 200 kPa) and noxious (> 200 kPa) mechanical stimuli (blue bars, Fig. 1B). By contrast, larvae transheterozygous for a viable combination of *InR* alleles (Im et al. 2018; Tatar et al. 2001) showed normal mechanical responsiveness at the low end of the noxious range (462 kPa or below) but increasingly impaired responsiveness at higher pressures (red bars, Figure 1B).

**Figure 1.**
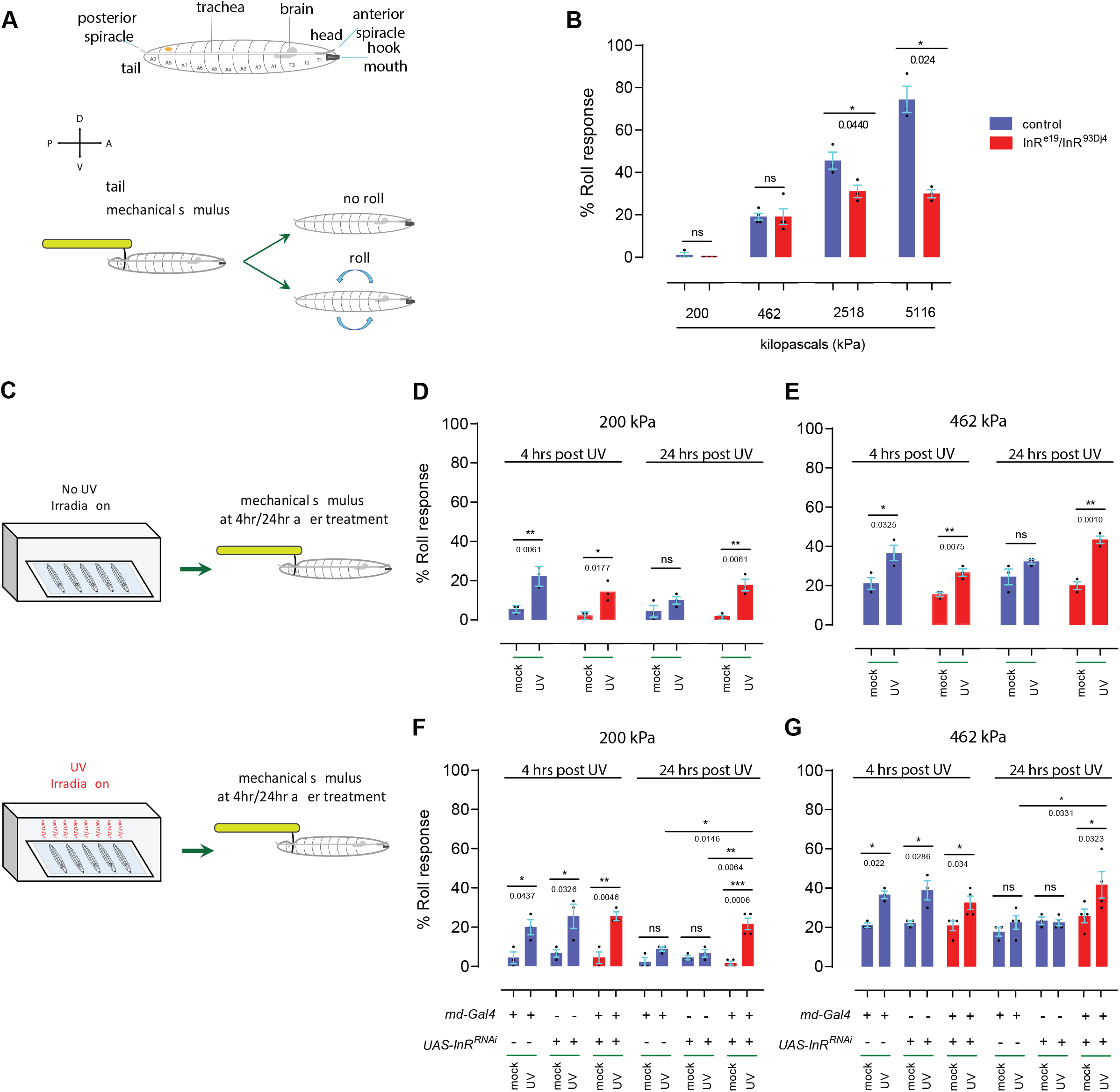
Mechanical hypersisitivity in flies lacking InR. (A)Schematic of mechanical nociception assay. Third instar Drosophila larvae (prominent anatomical features noted) are poked with a custom-designed and built larval Von Frey filament (see methods) exerting a defined amount of pressure. The resulting behavior is recorded. Sub-threshold pressures do not result in aversive rolling, while higher noxious pressures increase the percentage of larvae that perform the rolling behavior. (B) Behavioral dose-response curve of control (blue bar color) and InR mutant larvae (red bar color) of the indicated allelic combination. (C) Schematic of mechanical hypersensitivity assay. Larvae are either mock-irradiated (control) or UV-irradiated to induce epidermal tissue damage. At the indicated times post-irradiation, larvae were behaviorally tested using the mechanical nociception assay described in (A). (D-E) Quantitation of mechanical allodynia (D, mostly sub-threshold 200 kPa probe) and mechanical hyperalgesia (E, mildly noxious 462 kPa probe) in control (blue bar color) and InR mutant larvae (red bar color). (F-G) Quantitation of mechanical allodynia (F, mostly sub-threshold 200 kPa probe) and mechanical hyperalgesia (G, mildly noxious 462 kPa probe) in larvae with various genetic controls (*md-Gal4* alone, *UAS-InR^RNAi^* alone, blue color bars) or in larvae with InR targeted by *UAS-InR^RNAi^* in nociceptive sensory neurons (*md-Gal4* driver). Red bars indicate experimental larvae harboring both components of the Gal4/UAS system and thus expressing *UAS-InR^RNAi^* within nociceptive sensory neurons. For the quantitation panels, (B, D-G) each black dot and error bars indicate mean ± SEM with 3 to 4 sets of n = 30 larvae per timepoint and experimental condition.

Based on these baseline data we decided to use a 200 kPa probe (almost no aversive rolling in controls and mutants) to test for onset and persistence of injury-induced mechanical allodynia and a 462 kPa probe (~ 20% aversive rolling in controls and mutants) to test for induction and persistence of injury-induced mechanical hyperalgesia (See schematic, Figure 1C). Previously published experiments established that 4 hours post UV-irradiation is appropriate to test for onset of acute mechanical hypersensitivity and 24 hours is suitable to test for persistence and/or resolution of this acute response (Lopez-Bellido and Galko 2020). Although there were slight, non-significant variations in baseline responsiveness without UV irradiation (probably a function of *InR* mutant larvae being slightly smaller than controls), both control larvae and *InR* transheterozygotes showed newly acquired responsiveness (mechanical allodynia) to the 200 kPa probe and increased responsiveness (mechanical hyperalgesia) to the 462 kPa probe (Fig. 1 D-E). In control larvae, this transient hypersensitivity had resolved close to baseline responsiveness by 24 hours post-irradiation (Figure 1 D-E). By contrast, *InR* transheterozygotes exhibited persistent mechanical allodynia and hyperalgesia-they did not return back to baseline (Figure 1 D-E).

Finally, we tested whether InR function in mechanical allodynia is localized to peripheral multidendritic neurons, which are responsible for detecting noxious mechanical stimuli (Hwang et al. 2007; Kim et al. 2012; Lopez-Bellido et al. 2019b). Control larvae bearing only a Gal4 driver specific for nociceptive multidendritic sensory neurons (*md-Gal4*) (Gao et al. 1999) or the *UAS-InR^RNAi^* transgene (Ni et al. 2011) exhibited normal mechanical baselines and normal sensitization following UV-induced tissue injury (Figure 1 F-G). Larvae bearing both transgenes (and thus expressing *UAS-InR^RNAi^* in multidendritic nociceptive sensory neurons) also exhibited a normal baseline (~ 0 % responders to 200 kPa and ~ 20 % responders to 462 kPa) in the absence of injury and a normal peak of sensitization 4 hours post injury (Figure 1 F-G). This sensitization persisted in these larvae beyond the normal duration-the acute sensitization response did not resolve back to baseline at 24 hours post injury (Figure 1 F-G). These data indicate that in fly larvae InR is required for optimal baseline sensitivity to highly noxious mechanical stimuli and, similar to what has been observed for thermal sensitization (Im et al. 2018), also regulates the persistence of mechanical sensitization to both sub-noxious and mildly noxioius mechanical pressures.

### Sensory neuron specific deletion of mouse *Insr*

We wanted to test whether the abnormally persistent nociceptive sensitization observed in fly *InR* mutants might also be observed in a parallel experiment in mice. To do this, we bred mice heterozygous for a Cre driver (*NaV1.8-Cre*) specific for nociceptive sensory neurons (Agarwal et al. 2004) and homozygous for a floxed allele of *Insr* (Bruning et al. 1998), the locus that encodes the mouse insulin receptor. Using PCR primers specific for the intact floxed allele we detected an appropriately sized DNA band/product in all mice bearing the allele (Figure 2A). Only mice bearing the *NaV1.8-Cre* driver generated a PCR band specific for the *Cre* gene (also Figure 2A). Importantly, deletion of exon 4 of the *Insr* locus was tissue-specific-DNA harvested from DRG which contains the cell bodies of nociceptive sensory neurons, showed a PCR product specific for the excised locus when the Cre protein was present (Figure 2A). As expected, such a band was absent when skin tissue was used as the DNA source, whether or not the Cre transgene was present (Figure 2A). These data indicated that combining the *NaV1.8-Cre* transgene with the floxed allele of insr leads to a tissue-specific deletion of a portion of the mouse *Insr* locus.

**Figure 2.**
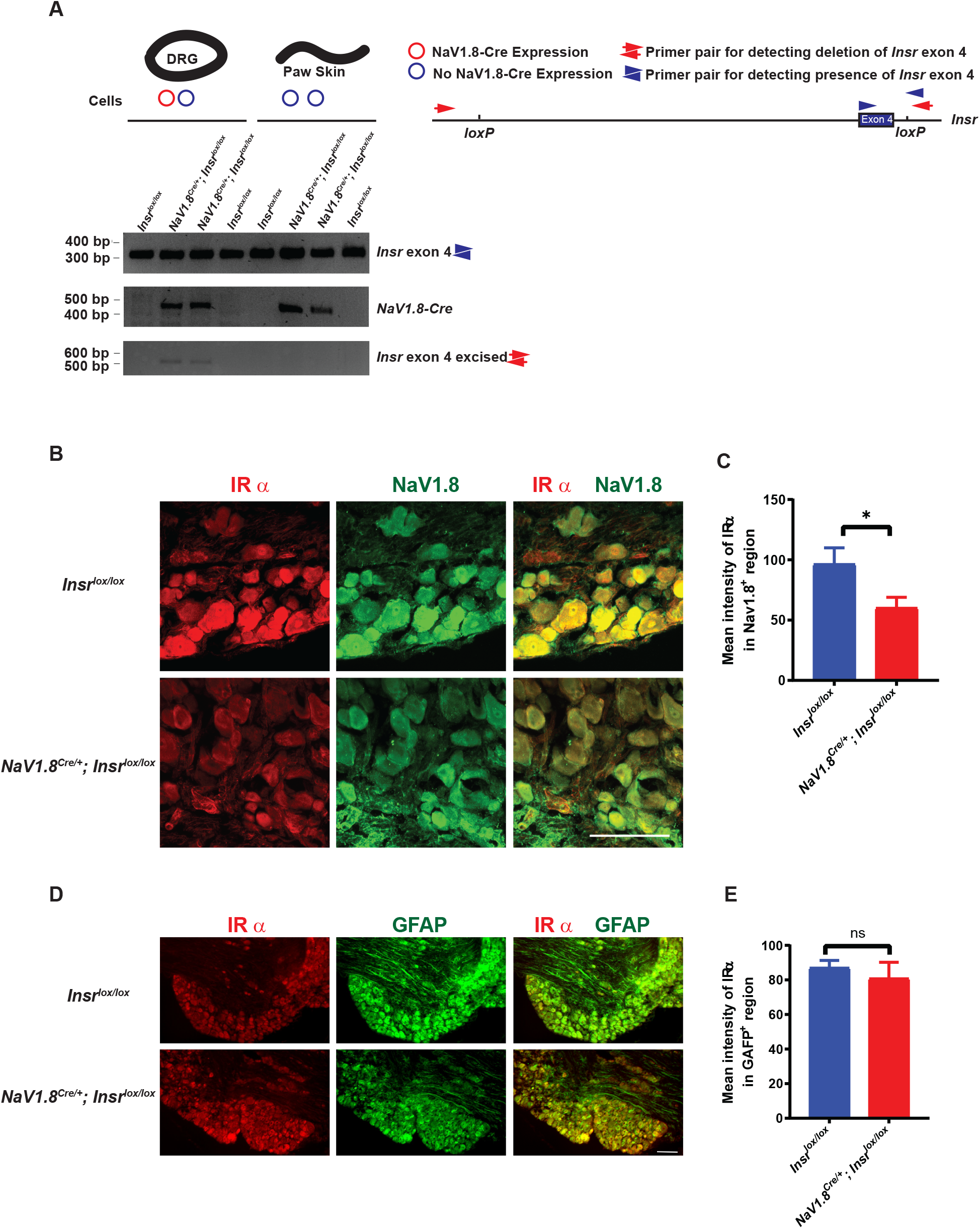
Sensory neuron specific deletion of mouse Insr. (A) Confirmation of Cre recombinase-mediated excision of insulin receptor exon 4 flanked by *loxP* sites-partial map of *Insr* locus show primer pair locations. Genomic DNA as template from DRGs (thick black circle) or paw skin (black wavy line) from mice of the indicated genotypes was PCR-amplified using primers suitable for detecting the unrecombined floxed *Insr* exon 4 (top row, primers = blue arrowheads), Cre recombinase (middle row, primers not shown), and deleted *Insr* exon 4 (bottom row, primers = red arrows). (B) Cryosections of DRG tissue dissected from *Insr^lox/lox^* and *NaV1.8^Cre/+^*;*Insr^lox/lox^* mice immunostained for Insulin receptor alpha subunit (IRα, red) and NaV1.8 (green). Scale bar, 100 μm. (C). Quantification of the intensity of IRα staining in NaV1.8+ DRG region of *Insr^lox/lox^* and *NaV1.8^Cre/+^*;*Insr^lox/lox^* mice. Insrlox/lox, n = 10 DRGs from 5 mice (2 females and 3 males). *NaV1.8^Cre/+^*;*Insr^lox/lox^*, n = 9 DRGs from 5 mice (2 females and 3 males). Mean ± S.E.M. Two-tailed unpaired Student’s t test was used for statistical analysis. *, p<0.05. (D) Cryosections of DRG tissue dissected from *Insr^lox/lox^* and *NaV1.8^Cre/+^*;*Insr^lox/lox^* mice immunostained for IR (red) and GFAP (green). Scale bar, 100 μm. (E). Quantitation of the intensity of IRα staining in GFAP+ DRG region of *Insr^lox/lox^* and *NaV1.8^Cre/+^*;*Insr^lox/lox^* mice. *Insr^lox/lox^*, n = 10 DRGs from 5 mice (2 females and 3 males). *NaV1.8^Cre/+^*;*Insr^lox/lox^*, n = 9 DRGs from 5 mice (2 females and 3 males). Mean ± S.E.M. Two-tailed unpaired Student’s t test was used for statistical analysis. ns, not significant.

We also examined insulin receptor protein levels in the DRG of mice harboring *Insr^lox/lox^* with or without the Cre driver (Figure 2B). We performed double immunofluorescence analysis with an antibody that recognizes NaV1.8, to mark cells that should be expressing the Cre driver, and with an antibody that recognizes the alpha chain of the mouse insulin receptor (IRα, Figure 2B). Quantitation of IRα levels revealed a reduction in the level of Insr in NaV1.8+ cells in mice harboring the Cre driver but not in littermate controls harboring only the *Insr^lox/lox^* allele (Figure 2C). We also performed double immunofluorescence analysis on the DRGs with an antibody that recognizes Glial Fibrillary Acidic Protein (GFAP), to label gial cells that should not be expressing the Cre driver, and with the antibody that recognizes the alpha chain of the mouse insulin receptor (Figure 2D). Quantitation of IRα levels revealed no significant difference in the level of Insulin receptor protein in GFAP+ cells between mice harboring the Cre driver and sibling controls with only the *Insr^lox/lox^* allele (Figure 2E). These data reveal that in addition to the DNA deletion at the *Insr* locus there is a reduction in IRα protein levels specifically in targeted NaV1.8+ sensory neurons.

### Mouse *Insr* regulates the persistence of inflammatory mechanical hypersensitivity in females

We next analyzed thermal and mechanical nociceptive sensitivity in the mice lacking *Insr* in NaV1.8+ nociceptive sensory neurons and relevant controls. Previously, it had been reported that mice with a broader deletion of *Insr* in sensory neurons using *Advillin-Cre* (Zurborg et al. 2011) had normal baselines for thermal and mechanical stimuli (Grote et al. 2018). Since flies lacking *InR* have normal baseline sensitivity but also exhibit persistent thermal (Im et al. 2018) and mechanical (Figure 1) hypersensitivity following injury, we decided to test mice following a model of inflammatory pain induced by intraplantar injection of CFA (see methods) that is conceptually similar to the UV damage assay employed in flies. Mice lacking *Insr* in NaV1.8+ neurons had normal thermal and mechanical sensitivity under baseline conditions, and no differences in the onset and resolution of thermal hyperalgesia following CFA injection into the hindpaw compared to controls (Figure 3A). Male and female mice lacking *Insr* in NaV1.8+ neurons showed normal baseline nociception and a normal onset of mechanical hypersensitivity (Figure 3B). However, in contrast with controls, the female mice showed a delayed resolution at later timepoints (days 14-21 following injury)(Figure 3B). Male mice of both controls and lacking *Insr* in NaV1.8+ neurons were not distinguishable from controls (Figure 3B). These data indicate that in an inflammatory pain assay mice lacking *Insr* in NaV1.8+ sensory neurons show a specific persistence of mechanical hypersensitivity only in females.

**Figure 3.**
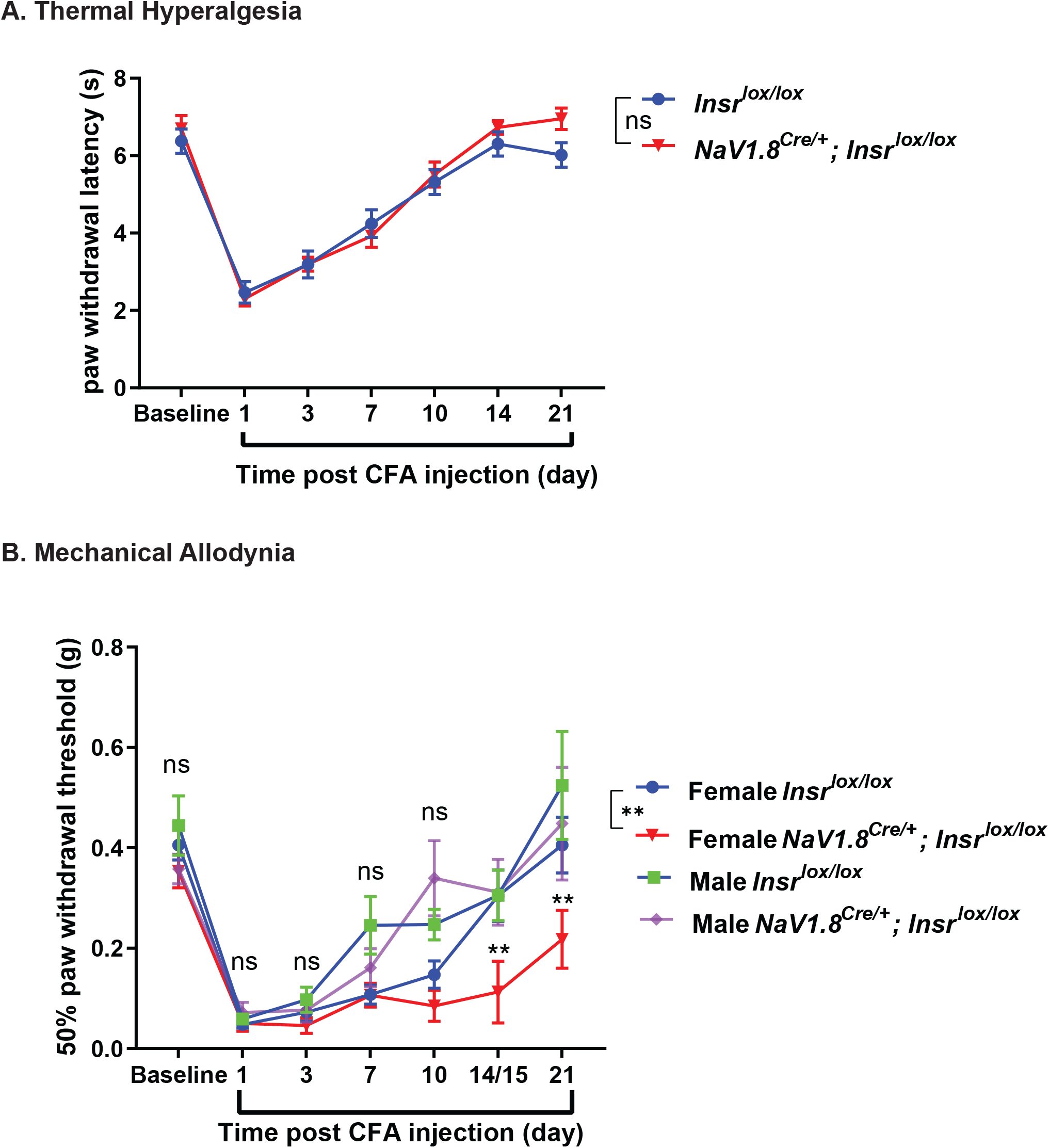
Behavioral analysis of mice lacking sensory neuron-specific Insr. (A) Quantitation of baseline thermal nociception and thermal hyperalgesia post CFA-induced local inflammation in *Insr^lox/lox^* controls and *NaV1.8^Cre/+^*;*Insr^lox/lox^* mice. *Insr^lox/lox^*, n = 8 (4 females and 4 males); *NaV18.^Cre/+^*;*Insr^lox/lox^*, n = 8 (4 females and 4 males). Mean ± S.E.M. Two-way repeated measures ANOVA followed by Sidak’s multiple comparison test was used for statistical analysis. ns, not significant. (B) Graph shows 50% paw withdrawal threshold of *Insr^lox/lox^* and *NaV1.8^Cre/+^*;*Insr^lox/lox^* mice with the mice separated by sex. *Insr^lox/lox^*, n = 24 (19 females and 5 males); *NaV1.8^Cre/+^*;*Insr^lox/lox^*, n = 19 (13 females and 6 males). *NaV1.8^Cre/+^*;*Insr^lox/lox^* mice of both sexes did not show defects in baseline mechanical nociception but female mice of this genotype exhibited a slow recovery in mechanical allodynia at 14/15 days post CFA-induced local inflammation. Mean ± S.E.M. Two-way repeated measures ANOVA followed by Tukey’s multiple comparison test was used for statistical analysis. ns, not significant. **, p<0.01.

### Metabolic activity in mice with a NaV1.8+ sensory neuron specific deletion of *Insr*

Although mouse sensory neurons are not generally thought to be a metabolic control tissue, we tested whether deletion of *Insr* in NaV1.8+ neurons had any systemic metabolic effects that might explain the observed persistent hypersensitivity in females. No sex-specific or other differences were observed between mutants and controls in body weight (Figure 4A), fasting blood glucose (a measure of baseline blood glucose level at a single point in time - Figure 4B), Hemoglobin A1C levels (a measurement of average blood glucose levels over the past two to three months- Figure 4C), and glucose tolerance (a measurement of how quickly fasted mice move new sugar from the blood into the tissues-Figure 4D). The lack of difference between controls and mutant mice lacking *Insr* in NaV1.8+ sensory neurons suggests that the observed delay in resolution of mechanical hypersensitivity in female mice (Figure 3) cannot be explained by systemic metabolic effects.

**Figure 4.**
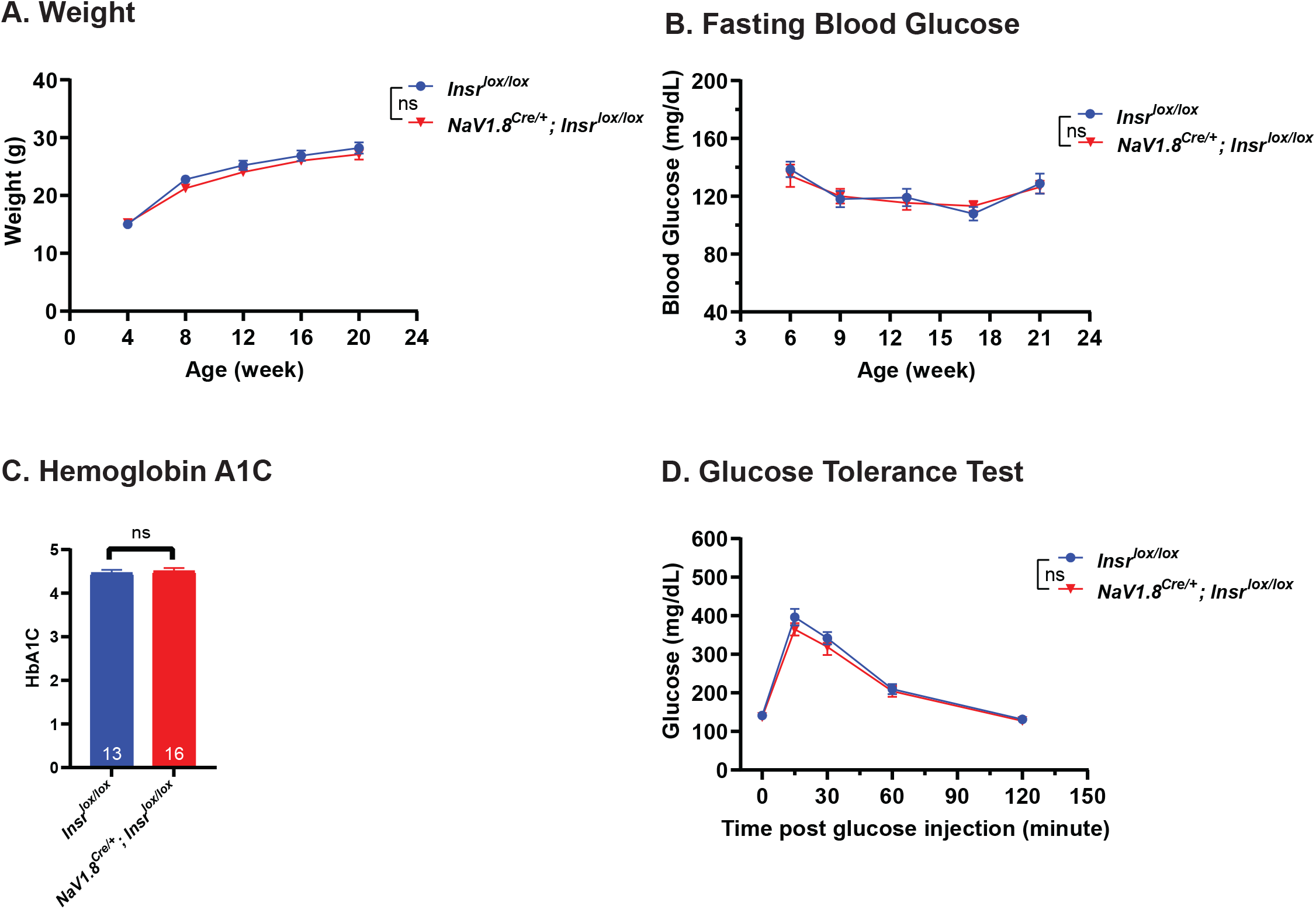
Metabolic analysis of mice lacking sensory neuron-specific Insr. (A). Body weight of *NaV1.8^Cre/+^*;*Insr^lox/lox^* mice and *Insr^lox/lox^* control mice as a function of the indicagted ages. *Insr^lox/lox^*, n = 22 (9 females and 13 males); *NaV1.8^Cre/+^*;*Insr^lox/lox^*, n = 21 (15 females and 6 males). Because no sex-specific differences were observed in this and the other metabolic measures (B-D) the data are not separated by sex. Mean ± S.E.M. Two-way repeated measures ANOVA followed by Sidak’s multiple comparison test was used for statistical analysis. ns, not significant. (B) Blood glucose levels of *NaV1.8^Cre/+^*;*Insr^lox/lox^* and *Insr^lox/lox^* mice as a function of the indicated ages. *Insr^lox/lox^*, n = 11 (5 females and 6 males); *NaV1.8^Cre/+^*;*Insr^lox/lox^*, n = 14 (8 females and 6 males). Mean ± S.E.M. Two-way repeated measures ANOVA followed by Sidak’s multiple comparison test was used for statistical analysis. ns, not significant. (C) Hemoglobin A1C levels of *NaV1.8^Cre/+^*;*Insr^lox/lox^* and Insr^lox/lox^ mice. Hemoglobin A1C levels were measured at 21 weeks of age. *Insr^lox/lox^*, n = 13 (6 females and 7 males); *NaV1.8^Cre/+^*;*Insr^lox/lox^*, n = 16 (9 females and 7 males). Mean ± S.E.M. Two-tailed unpaired Student’s t test was used for statistical analysis. ns, not significant. (D) Glucose tolerance in *NaV1.8^Cre/+^*;*Insr^lox/lox^* and *Insr^lox/lox^* mice. An IPGTT was used to measure glucose levels immediately prior to glucose stimulation and at the indicated times thereafter. *Insr^lox/lox^*, n = 13 (6 females and 7 males); *NaV1.8^Cre/+^*;*Insr^lox/lox^*, n = 16 (9 females and 7 males). Mean ± S.E.M. Two-way repeated measures ANOVA followed by Sidak’s multiple comparison test was used for statistical analysis. ns, not significant.

## Discussion

A prior study from our lab established that *Drosophila InR* regulates the persistence of injury-induced thermal nociceptive hypersensitivity (Im et al. 2018). We report here that *Drosophila* larvae mutant for *InR* or expressing *UAS-InR^RNAi^* transgenes in nociceptive sensory neurons exhibit normal baselines for mechanical nociception and normal acute sensitization. However, these larvae exhibit persistent mechanical allodynia and hyperalgesia following UV injury. This suggests that, in flies, InR regulation of the persistence of injury-induced nociceptive sensitization is not specific to heat sensitivity. This is perhaps not surprising since *Drosophila* class IV multidendritic (md) neurons are multimodal, mediating nociceptive responses both to high-threshold thermal stimuli (Babcock et al. 2009; Tracey et al. 2003) and mechanical stimuli (Hwang et al. 2007; Kim et al. 2012).

These *Drosophila* results prompted us to investigate whether *Insr* similarly regulates the persistence of inflammatory nociceptive sensitization in mice. We combined a floxed allele of *Insr* (Bruning et al. 1998) with a Cre driver specific for nociceptive sensory neurons (Agarwal et al. 2004) to delete *Insr* in these cells. As reported with a deletion of *Insr* in a broader subset of sensory neurons and in gut cells using *Advillin-Cre* (Zurborg et al. 2011), thermal and mechanical nociceptive baselines were normal in these mice (Grote et al. 2018). Here, since we also did not detect differences in baseline mechanical or thermal sensitivity, we also tested nociceptive sensitization following inflammatory injury. Baseline nociception for both sensory modalities was normal and acute thermal and mechanical hypersensitivity peaked normally in mice lacking *Insr* in NaV1.8+ sensory neurons versus controls. Two aspects of the response were different, compared to the results in *Drosophila*. First, no persistent thermal nociceptive hypersensitivity was observed. Second, the persistent mechanical nociceptive hypersensitivity was specific to female mice. The observation of mechanical hypersensitivity is consistent with reports that *NaV1.8-Cre* expresses in at least a subset of mechanically responsive sensory neurons (Daou et al. 2016; Shields et al. 2012). Several possibilities may explain the differences between flies and mice. First, the architecture of nociceptive sensory neurons in mice is more diverse, with a more clear physical and functional separation between neurons mediating thermal and mechanical responses (Jenkins and Lumpkin 2017; Le Pichon and Chesler 2014). Second, there have been reports of differing nociceptive sensitivity between male and female diabetic patients (Ruau et al. 2012).

One other study (Grote et al. 2018) has reported on sensory phenotypes of mice with a deletion of *Insr* in nociceptive sensory neurons. The driver used in this former study was *Advillin-Cre*, which expresses in a broader subset of sensory neurons but also expresses, like the *Advillin* promoter in gut endocrine cells and enteric neurons (Hunter et al. 2018; Melo et al. 2020). In this study normal thermal and mechanical nociceptive baselines (without injury) were observed. This study did not observe a difference in blood sugar levels but did observe differences in serum insulin levels and glucose tolerance that appeared in older mice (29+ weeks of age). Some of those same measures appeared normal in the experiments reported here. This raises the possibility that the metabolic effects observed previously were mediated either by sensory neurons not targeted by the *NaV1.8-Cre* driver used here or by enteric neurons or gut endocrine cells that express *Advillin*. We view the latter as more likely given that the gut is a known metabolic regulatory tissue (Drucker 2007). Further experimentation will be needed to parse out these differences.

Finally, this study may have implications for the etiology of diabetes-associated pain (Obrosova 2009). Some have argued that neuropathic pain in diabetic patients may have a predominantly cental nervous system origin (Fischer and Waxman 2010) whereas others have suggested possible roles for peripheral insulin signaling (Grote and Wright 2016). In *Drosophila*, however, the persistent thermal nociceptive sensitization phenotype seen with sensory neuron specific knockdown of InR is precisely phenocopied in models of type 1 and type 2 diabetes (Im et al. 2018). Further, a recent study suggests that mechanical hypersensitivity is also observed in the type 2 diabetes model in fly larvae (Dabbara et al. 2021). In diabetic patients, correlation of blood sugar control with pain hypersensitivity has been difficult to establish (Chan et al. 1990). Prominent models for diabetes-associated hypersensitivity include a secondary consequence of poor vascular tone (Powell et al. 1985) and possible adverse affects of glycation byproducts on sensory neurons (Orestes et al. 2013). Our results suggest that another model is worth considering and testing further-that mechanical hypersensitivity may be related to poor insulin signaling directly within nociceptive sensory neurons, particularly in female vertebrates.

## Acknowledgements

*Drosophila* stocks were obtained from the Bloomington *Drosophila* Stock Center (Bloomington, IN) and the Vienna *Drosophila* RNAi Center (Vienna, Austria). We thank Ashok Subedi for assistance with DRG dissection. This study was supported through R21NS126929 to MJG and AK. Veterinary resources were supported by NIH grant CA16672. Dr. Richard Behringer and Galko laboratory members read and commented on the manuscript.

